# Deriving the cone fundamentals: a subspace intersection method

**DOI:** 10.1101/2024.02.04.577470

**Authors:** Brian A. Wandell, Thomas Goossens, David H. Brainard

**Affiliations:** Psychology Department, Stanford University, Stanford, CA, 94305 USA; Psychology Department, University of Pennsylvania, Philadelphia, 19104 USA

## Abstract

Two ideas, proposed by Thomas Young and James Clerk Maxwell, form the foundations of color science: (1) Three types of retinal receptors encode light under daytime conditions, and (2) color matching experiments establish the critical spectral properties of this encoding. Experimental quantification of these ideas are used in international color standards. But, for many years the field did not reach consensus on the spectral properties of the biological substrate of color matching: the sensitivity of the *in situ cones* (cone fundamentals). By combining auxiliary data (thresholds, inert pigment analyses), complex calculations, and color matching from genetically analyzed dichromats, the human cone fundamentals have now been standardized.

Here we describe a new computational method to estimate the cone fundamentals using only color matching from dichromatic observers. We show that it is not necessary to include data from trichromatic observers in the analysis or to know the primary lights used in the matching experiments. Remarkably, it is even possible to estimate the fundamentals by combining data from experiments using different, unknown primaries. We then suggest how the new method may be applied to color management in modern image systems.

## 1. Introduction

> « *It appears therefore that the result of any mixture of colours, however complicated, may be defined by its relation to a certain small number of well-known colours. Having selected our standard colours, and determined the relations of a given colour to these, we have defined that colour completely as to its appearance. Any colour which has the same relation to the standard colours, will be identical in appearance, though its optical constitution, as revealed by the prism, may be very different. »*

James Clerk Maxwell [1]

More than two centuries ago artisans of color prints in both Europe and Asia knew that a wide range of color appearance could be achieved by the mixture of three primary colors [2]. Thomas Young explained the necessity and sufficiency of mixing three primaries by the idea that the retina contains three types of light sensitive cells [3]. Biology, not physics, was his explanation of trichromacy.

More than 150 years ago, James Clerk Maxwell developed methods to quantify the biological encoding of color [4–8]. Critical to the present paper, Maxwell implemented an apparatus to measure what we now call the color matching functions. Subjects adjusted the intensity of triplets of spectral lights to match a constant daylight. The experiment is linear [9], and this enabled Maxwell to derive the spectral color matching functions through clever experimental choices and simple algebraic manipulation (Figure 1). He tabulated measurements from two typical subjects (trichromats), his wife and himself^1^. As the opening quotation shows, Maxwell recognized that the mixture of three spectral lights is sufficient to match a broadband light. His measurements guided the first implementation of color photography by Sutton and Maxwell [10].

**Fig. 1.**
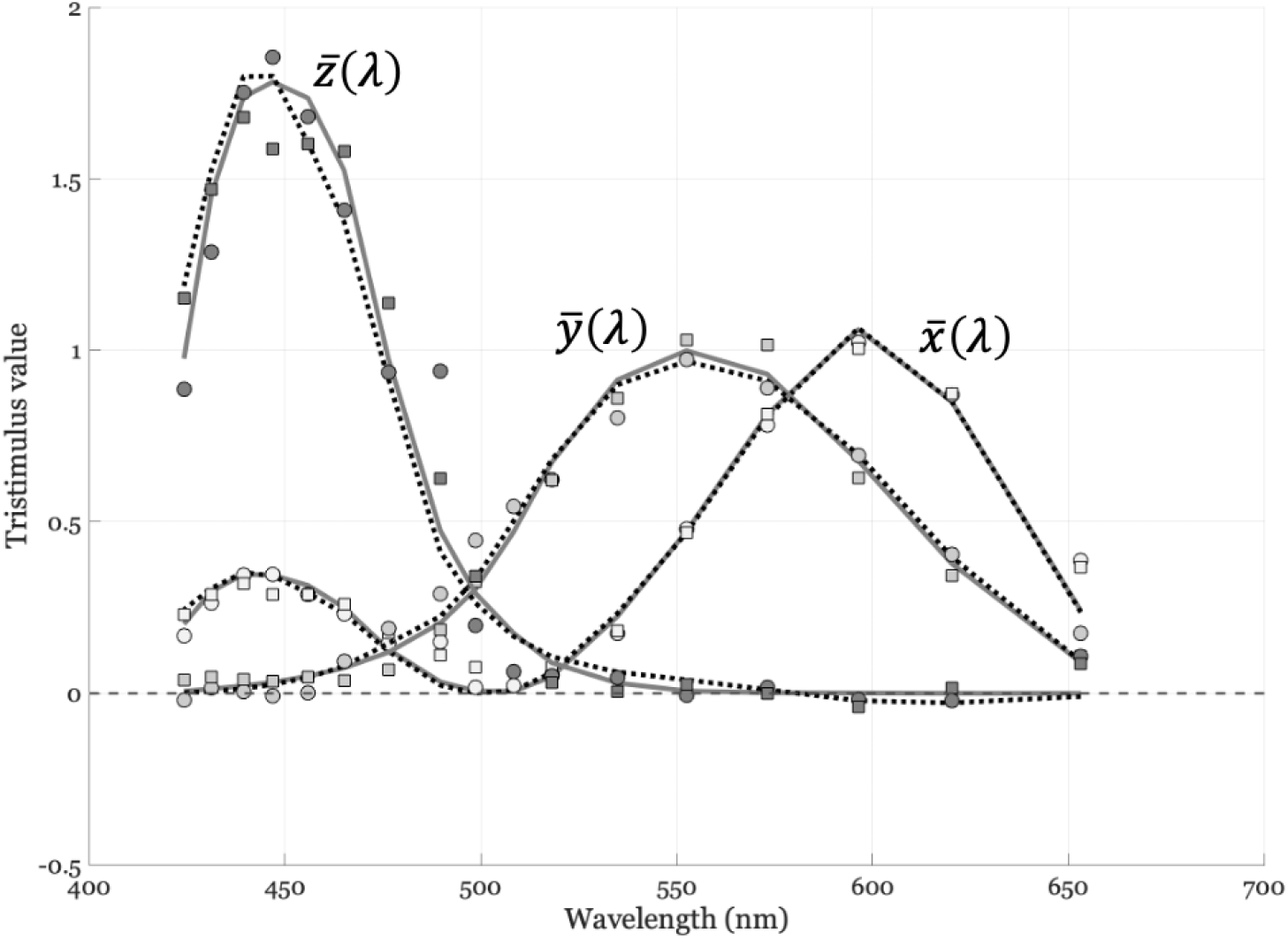
The color matching data reported by Maxwell [1]. The solid curves are the CIE (1931) tristimulus color matching functions. The dashed curves are the current CIE cone fundamentals [11, 12] linearly transformed to match the CIE 1931 functions. The points are color matching functions measured by Maxwell in two observers (J,squares; K,circles), also linearly transformed to match the CIE 1931 functions. The general agreement between Maxwell’s data and the CIE curves was pointed out by Judd, who provided a conversion of Maxwell’s wavelength units (Paris inches) to nanometers [6]. Here we include a small (10 nm shift) correction to Judd’s estimate of the Maxwell’s wavelengths, as well as show the comparison with the current CIE cone fundamentals. The data and calculations to produce this figure are included in the software repository associated with this paper (see fig01Maxwell/maxwellCMF2CIE.m)

At the time Maxwell made his measurements, Young’s hypothesis was not universally accepted. The distinguished physicist, Brewster, explained color as a physical property of light. Helmholtz initially believed trichromacy to be in error, before reversing his view [13–16]. Maxwell swept these concerns aside by experimental demonstration. He observed that interposing a color filter between the eye and a pair of lights that look the same could break the match; this would not happen if Brewster was correct and the lights were physically identical.

But what are the the spectral sensitivities of the biological substrate suggested by Young? Such sensitivities are now referred to as the cone fundamentals. Since experimental color matching functions depend on the spectral lights used as primaries, they cannot be taken directly as estimates of the cone fundamentals. Based on the linearity of the color matching experiment, however, it can be shown that the cone fundamentals are a linear transformation away from any given set of spectral color matching functions [17]. Thus estimating the cone fundamentals can be framed as determining the appropriate linear transformation.

Maxwell initiated color matching studies with dichromats [4] before moving on to the problem of unifying electricity and magnetism. It was Helmholtz’ student, König, who realized that if the cone fundamentals of dichromats are a subset of those of trichromats (reduction dichromats), it should be possible to use the color confusions of dichromats to infer the cone fundamentals. König and Dieterici developed a method for measuring the dichromatic color confusion lines to estimate the cone fundamentals [18]. The approach they implemented had dichromats set trichromatic matches. The data and theory has been formalized and explored by many others [19–24].

A project to marshal modern methods to establish the cone fundamentals culminated in the adoption of an excellent and secure standard [25]. This work built on prior analyses [23, 26, 27], and it incorporated auxiliary data including threshold measurements from dichromats, genetic profiling to assure the reduction hypothesis, and estimates of the inert pigments (lens, macula) of the eye in individual subjects. The methods, and their relationship to the careful work of many earlier investigators, are explained in Stockman’s authoritative review [12]. The cone fundamentals as well as other fundamental colorimetric data are readily available in tabulated form [28].

The result we report here is a new computational method for estimating the cone fundamentals directly from dichromatic color matching experiments, requiring only two primaries. We explain the method and how it differs from the classic approach introduced by König. We also provide an implementation in the software repository accompanying this article; at its core, the method requires only a few lines of code in a modern computational environment.

Because the idea underlying the new method is simple, it provides a useful way to introduce the color science to students. Because it is based on standard linear algebraic methods, it may find application in modern color image systems as well as in the adjacent fields of robotics and computer vision.

In the following, we explain the new method and its implementation. We then illustrate its application using data from F.H.G. Pitt, W.D. Wright and colleagues, collected in the 1930s and 1950s. It is particularly satisfying that the careful empirical work of these scientists is still relevant and reproducible. We outline how these methods may be applicable to image systems, robotics, and computer vision applications.

## 2. Cone fundamental estimation

Typical subjects have three types of cones, commonly referred to as long-wavelength (L), middle-wavelength (M), and short-wavelength (S) cones. Reduction dichromats are missing one type of cone, protanopes (no L), deuteranopes (no M), tritanopes (no S). The cone fundamentals for each subject are cornea-referred; the fundamentals combine the spectral properties of the cornea, lens, and macular pigment. These elements differ between people and the lens absorption increases with age [29–31]. Current standards for cone fundamentals account for these factors, and indeed attention to such variation is key to maximizing the accuracy of the standards. Here we focus on the underlying principles and not on applying them with the level of precision that would be required to create a standard. Thus, the illustrative analysis of dichromatic matches we present below are based population average data.

Color matching experiments can be described using linear algebra because the experiments satisfy the principle of additivity that defines a linear system [32]. Specifically, consider the functions that describe the spectral energy in two test lights, *e*_1_ (λ) and *e*_2_ (λ). In trichromats, the color matching experiments maps any test light spectral energy distributions to the intensities of three primaries. Additivity means that the match to the sum of the lights is the sum of the matches to each light alone,

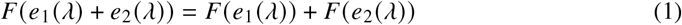

where *F* maps the spectral power distribution to three primary intensities. In color science, such additivity is referred to as Grassmann’s Laws [9]. Because the experiment is additive, there are three functions, *c*_*i*_ (λ), such that the inner product, ⟨*c*_*i*_, *e*_1_⟩, is the intensity of the *i*^*th*^ primary in the match [32]. The *c*_*i*_ (λ) are called the color matching functions.

The functions *e*_*i*_ and *c*_*i*_, can be discretized and represented as N-dimensional vectors, where *N* is equal to the sampling resolution of the wavelength input spectrum, say from 380 nm to 780 nm in 5 nm increments. The color matching functions, *c*_*i*_ (λ), can be combined into the columns of a matrix **C** ∈ ℝ^*N*×3^. By linearity, we can calculate the intensities of the primaries that match a test light, **C**^*t*^**e**_*i*_. This formulation achieves Maxwell’s goal: we use the color matching functions to calculate the intensities of three primary lights that would match any test light. If two lights, presented in the same context, are matched by the same primaries, **C**^*t*^**e**_1_ = **C**^*t*^**e**_2_, the lights will appear the same. The linearity principle and the linear algebraic formulation are fundamental to many fields of science and engineering [33].

### 2.1. The color matching functions

Measurements made using different primary lights yield different color matching functions. Any pair of color matching functions will be related to one another by a 3 × 3 linear transformation [17]. Because the cone fundamentals can serve as a set of color matching functions, measured color matching functions must be within a 3 × 3 linear transformation of the cone fundamentals.

Suppose we represent the cone fundamentals as *l* (λ), *m* (λ), *s* (λ), discretized in vector form as **l, m, s** ∈ ℝ^*N*×1^. Dichromats need only two primary lights to match any light. The color matching functions of a dichromat will be within a 2 × 2 linear transformation (**A**_*p*_, **A**_*d*_, and **A**_*t*_) of their two cone fundamentals. In matrix form we have

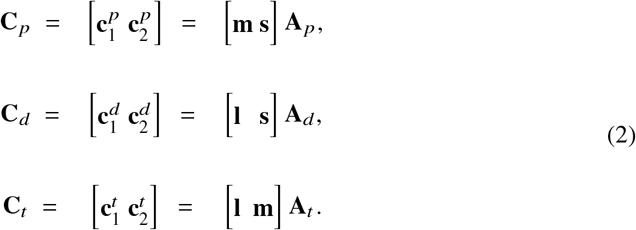

The dichromatic color matching matrices **C**_*p*_, **C**_*d*_, **C**_*t*_ ∈ ℝ^*N*×2^.

### 2.2. Cone fundamental estimation by subspace intersection

The three cone fundamentals can be estimated from the color matching functions of the three reduction dichromat pairings. The cone fundamental that is in common between each pair can be found by calculating the null space of the *N* × 4 matrix created by combining the dichromatic color matching functions.

For example, consider the color matching functions of a protanope and deuteranope. These two individuals share the M-cone fundamental **m**. Consequently, there is a linear combination of each of their color matching functions that equals the M-cone fundamental. When the coefficients of these linear combinations are represented as the vectors **x**_*p*_ ∈ ℝ^2^ and **x**_*t*_ ∈ ℝ^2^, we can formalize the above statement as:

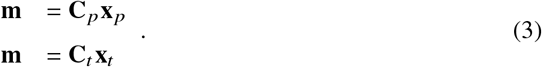

Subtracting the two equations we have

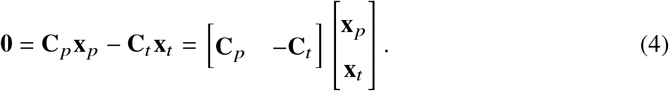

The space of all solutions (**x**_*p*_, **x**_*t*_) to Equation 4 is the null space of the matrix [**C**_*p*_ − **C**_*t*_ ] : the set of vectors that the matrix maps to zero. The null space can be calculated using the linear algebra packages in all major programming languages. In this case, the null space is one-dimensional, and it can be expressed as the vector times a scalar, 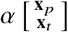. The cone fundamental in common to the two dichromats, **m**, can be estimated using the null space vector:

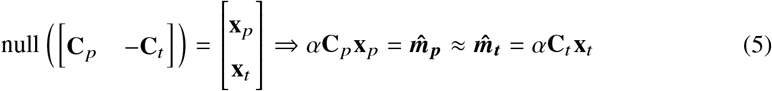

Equation 5 represents the result as an approximation because measurement noise and the possibility of individual differences in the inert pigments, photopigment absorbance spectra, and photopigment densities means there will not be a true null space, 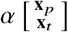. A consequence is that the two estimates, 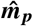 and 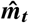, will differ from one another. Moreover, any solution will depend on the numerical algorithm we use to approximate the null space. Here, we estimated the null space using the Singular Value Decomposition (SVD), which returns the vector with the smallest mean squared error.

It is also possible to estimate the null space as an explicit numerical optimization, i.e. find the minimum of ∥**C**_*p*_**x**_*p*_ − **C**_*t*_ **x**_*t*_ ∥ subject to a constraint that prevents the degenerate solution **x**_*p*_ = **x**_*t*_ = 0. This approach would allow incorporation of other constraints, for example that the estimated cone fundamental be non-negative. We plot the estimate based on **x**_**p**_ in Figure 4. We implement other methods in the GitHub software repository^2^ and plot the different estimates.

### 2.3. Geometric interpretation

The calculation in Section 2.2 makes no assumption about the primary lights used to measure the color matching functions. In fact, the primaries used for the different dichromats can differ across the dichromats and need not be known.. To understand how this can be, a geometric interpretation may be helpful.

Each color matching function can be represented as a high-dimensional vector, with dimensionality equal to the number of sample wavelengths. The two color matching functions from each dichromat span a plane through the origin in this wavelength space. The two planes for any pair of dichromats both contain a line that represents their common cone fundamental. This line, which also passes through the origin, is found at the intersection of the two planes. It is this intersection that is calculated in Equation 5.

We can visualize this calculation for a low-dimensional example. Suppose the color matching functions are sampled at only three wavelengths, so that the vector representing each dichromatic color matching function is three-dimensional. Since color matching functions for a given observer are within a linear transform of each other, the two vectors define a plane comprising all possible color matching functions (Figure 2). The planes for the deuteranope and protanope both contain the vector representing the S-cone fundamental, so that the two planes intersect in the S-cone vector. No matter which primary lights were used to measure the color matching functions, the planes spanned by the color matching functions are the same. Hence, the intersection is independent of the primary lights.

**Fig. 2.**
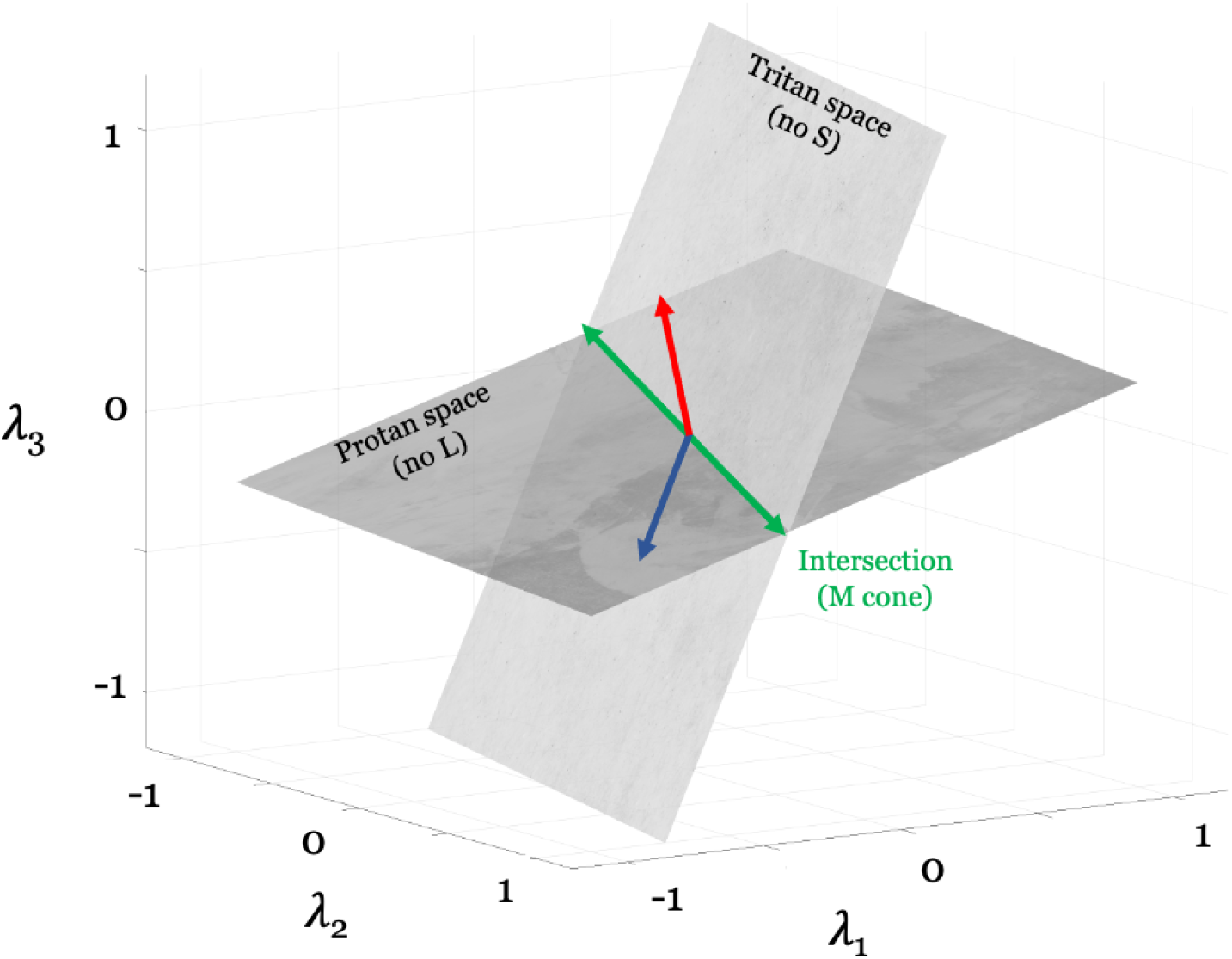
When sampled at only three wavelengths, the color matching functions are three-dimensional, and the pair of functions for each dichromat define a plane. The planes for a tritanope and a protanope are illustrated schematically. The plane for each dichromat contains the vectors defining the two cone fundamentals for that dichromat: the L- and M-cone fundamentals for the tritanope; the M- and S-cone fundamentals for the protanope. Note that the plane spanned by the color matching functions for each dichromat is independent of the primary lights used to measure them; this is because each color matching function is a linear transformation of the dichromat’s cone fundamentals and thus must lie in the plane spanned by those fundamentals. The intersection of the planes for the two dichromats must contain the cone fundamental they share, here the M-cone fundamental. Since the intersection of two planes is a line, this line defines the cone fundamental up to a scalar.

The analysis we describe for deriving cone fundamentals using data from reduction dichromats differs from the classic approach [20, 34], which is based on König’s important work. That approach, which is also included as a tutorial program in our software repository (see “tutorials/konig”), begins in the three-dimensional space defined by the trichromatic color matching functions, not the wavelength representation. In the trichromatic space, confusion lines are found for each type of dichromat. Differences between stimuli on a confusion line are invisible to the dichromat, defining a stimulus manipulation that isolates the dichromat’s missing cone class. With knowledge of the trichromatic directions that isolate each cone class, it is possible to recover the linear transformation between the trichromatic color matching functions and the cone fundamentals [35]. The interested reader is referred to the tutorial program for details. ^3^ The classic method requires dichromatic confusion measurements in a trichromatic color space. By analyzing the color matches directly in wavelength space, the subspace-intersection method does not rely on confusion lines; nor does it require making measurements with the same primary lights for different subjects.

## 3. Results

The subspace-intersection method to infer the cone fundamentals requires measurements of the color matching functions from all three types of dichromats. Analyses of dichromatic color-matching functions were initiated in the latter part of the 19^th^ century by König [18]. Color matching functions of dichromats were later reported in a series of papers from Wright and Pitt [36–38]. The accompanying software repository includes these data, comparisons between them and the modern standards, and some caveats about the data (see Figure 3).

**Fig. 3.**
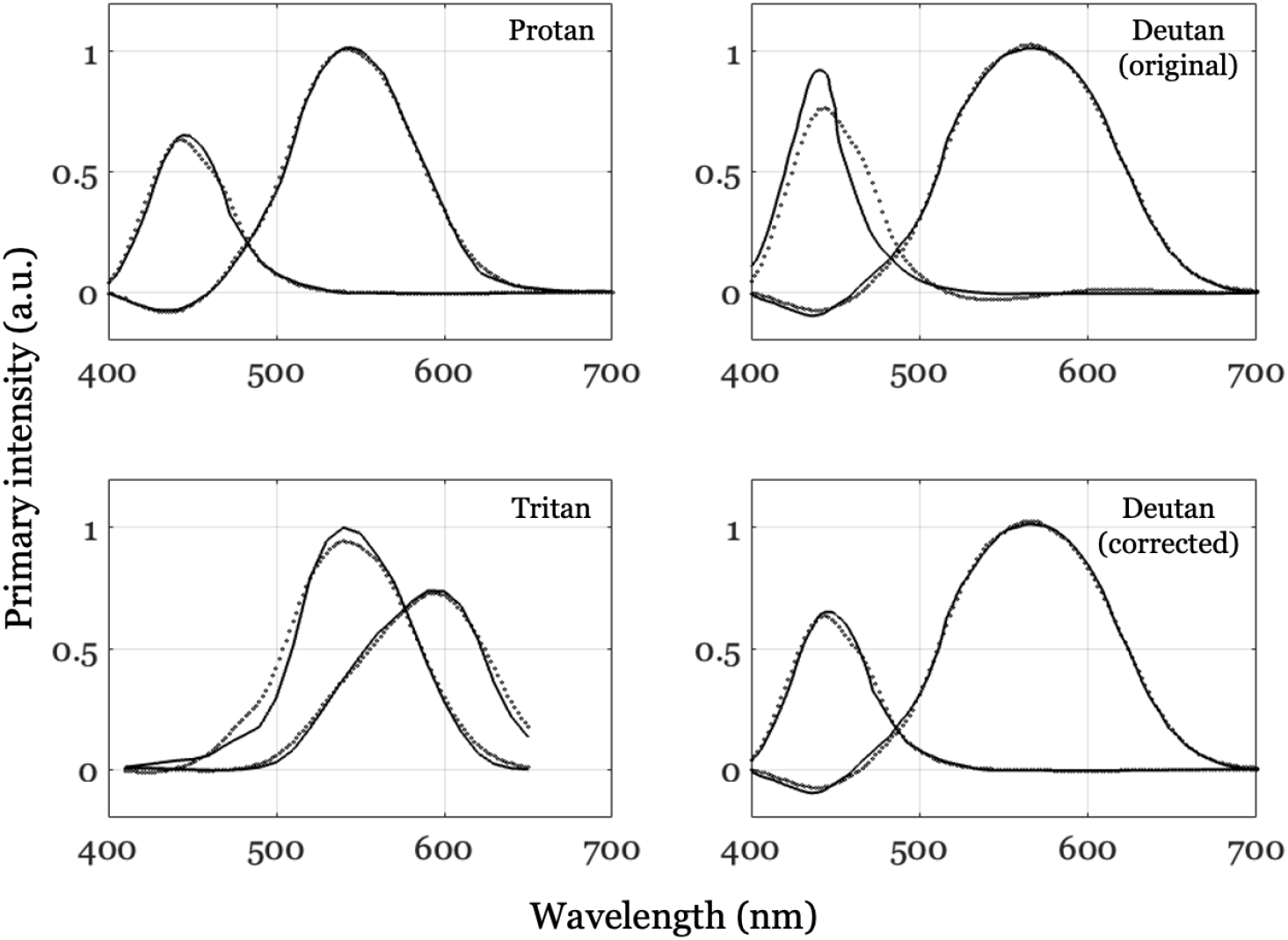
The Wright dichromatic color matching functions (solid) compared with a linear transformation of the CIE cone fundamentals (dotted). The protan and deutan functions were converted to digital form from figures in Wright’s book [37]. The tritanopic data were provided as tables [38]; notice that the wavelength range for the tritanopes is narrower. The original deutan data have an implausible short-wavelength color matching function (gray dashed curves). The estimates in Figure 4 were made substituting the protan blue primary for the deutan (corrected). For both of these dichromats the short-wavelength primary should be dominated by the common S-cone. This figure is rendered by the script fig03WDWDichromats/wdwStockman.m

**Fig. 4.**
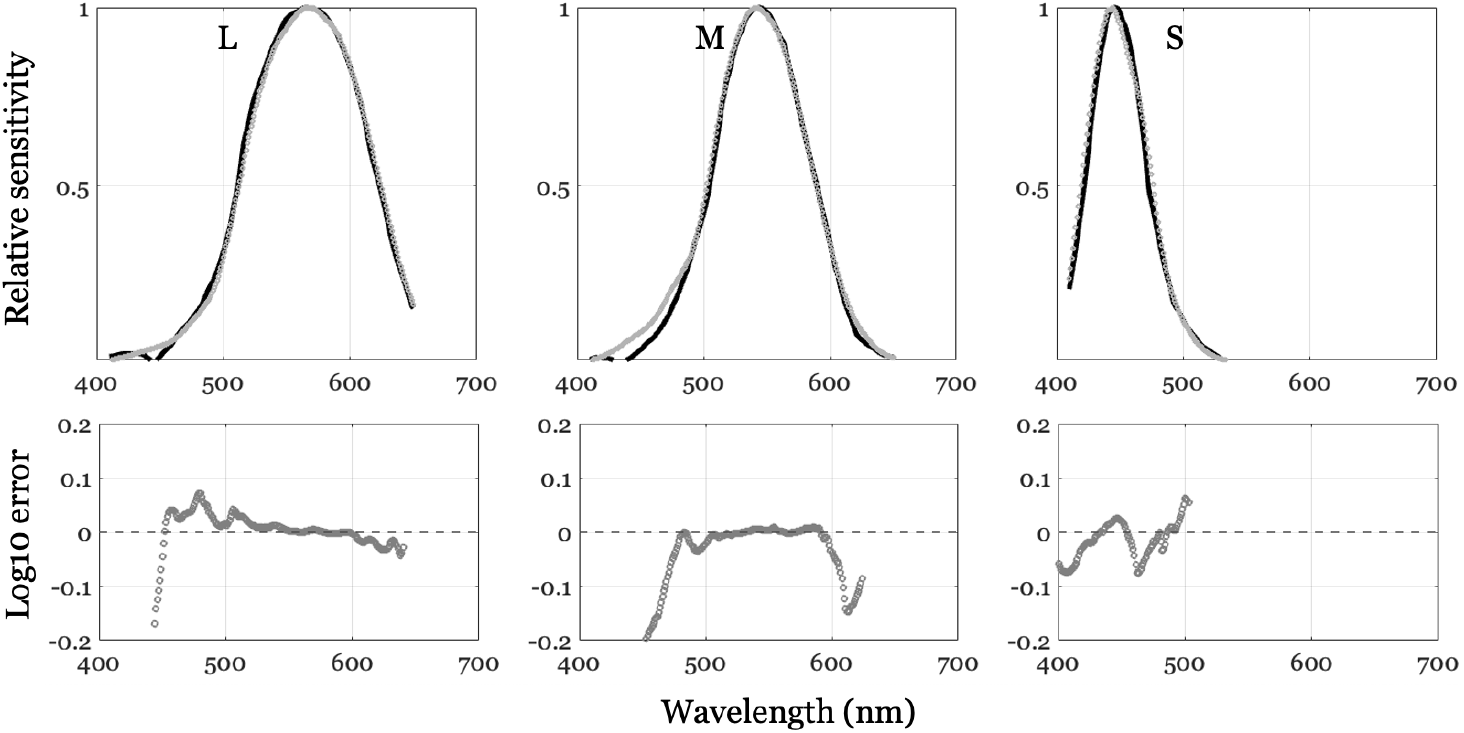
Cone fundamentals derived from the Wright and Pitt dichromatic color matching functions. The top row compares the better of the two estimated fundamentals (dark, **x**_*p*_ solution) with the CIE cone fundamentals (gray) [12]. The estimates are calculated from the modified Wright data (Figure 3). The bottom row shows the log10 difference between the curves for CIE values greater than 0.05 (peak = 1). The calculation and graphs are produced in the software repository script fig04ConeEstimates/s_cfConeEstimates.m. A second estimate is similar; it is plotted in the repository script.

Applying the methods from Section 2.2 to the dichromatic data of Pitt and Wright [36–38] yields estimates of the cone fundamentals (Figure 4). There are some limitations to the data, which were measured at different times and labs in experiments spanning more than a decade. (1) None of the subjects were genetically tested to show they are pure reduction dichromats, so it is likely that the mean values include measurements from dichromats who are not of the pure reduction type. (2) Like others [19], we believe there is an error in the curve drawn for the blue primary of the deutan subjects; we substituted the blue primary of the protan subjects because it should be very close to the same for both dichromats. With these limitations, the estimates based on three pairs of average dichromatic color matching functions and the current CIE cone fundamentals agree over most of the range to within about 0.1 log_10_ units. The main differences are in the M-cone fundamental below 500 nm, and the estimated L-cone fundamental at wavelengths below 475 nm.

## 4. Discussion

This paper describes data and theory developed over two centuries, making it impossible to offer a complete review of the relevant literature. We limit the discussion to a few points: (a) the reproducibility of the historical literature, (b) the value of knowing the cone fundamentals, (c) color appearance and color matching, and (d) virtual channels in image systems, robotics, and computer vision applications. We plan to explore the image systems applications more fully in a separate paper.

### 4.1. Reproducibility

Color matching measurements are in the great tradition of psychophysics; stimuli have precise physical units and experiments use simple and highly reliable psychological judgments. The tabulated data in the historical papers made it possible for us - and Judd before us - to compare Maxwell’s measurements acquired more than a century ago with the CIE standard color matching functions used by thousands of modern image systems engineers to design products viewed by billions of people. We also quantify the difference between the estimates with the modern cone fundamentals.

A small amount of the data needed for this paper was either not tabulated, represented with units or conventions that are no longer used, or presented in ways that required us to recover the measurement from a graphed quantity. These limitations introduced uncertainty in our estimates. It is both obvious and worth emphasizing that tabulated measurements in internationally accepted units provide a firmer basis for reproducibility. The psychophysical methods and tabulation of the data provide an excellent basis for reproducible research.

### 4.2. Biological substrate

The excellent quantitative agreement between behavioral color matching data and cone photocurrent action spectra [39] establishes a connection between perception and a neural substrate; it is one of the clearest connections in all of neuroscience. It is worth remembering that the CIE color matching standards were used in many applications before the neural substrate was established. The identification of the substrate is intellectually satisfying and adds value; but knowledge of the substrate was not critical to many successful applications. Had Hering been correct, and the cone photopigments encoded light using an opponent mechanism [40], Maxwell’s color matching functions would still have been useful for color photography.

An important value of the CIE cone fundamentals is that they provides a mechanistic description. Investigators analyzing the cone fundamentals needed to consider several components of the biological substrate: the lens, macular pigment, and photopigments. The contributions of these separate mechanisms is useful to understand and model population variance ([32], Fig 9; [30]) and to predict the consequences one might expect from certain diseases. The color matching functions are a very useful phenomenological model; cone fundamentals are a mechanistic model which is intellectually satisfying.

### 4.3. Color appearance

Color appearance is an important part of most peoples’ experience of the world; the topic has drawn the interest of scientists and philosophers over the centuries. It is striking, therefore, that there is almost nothing of color appearance in Maxwell’s original color matching experiment. His subjects matched only a white light. He chose this approach because the matches can be made with great precision, and he could do the rest by numerical calculations.

In the century following Maxwell’s work, prominent authors criticized the Young-Maxwell-Helmholtz color research program for failing to address color appearance^4^. We view this criticism as unfair because many color scientists - including Goethe, Helmholtz and Maxwell - did consider color appearance in the broader scope of their work.

It is worth considering that the progress from the Young-Maxwell-Helmholtz school was helped by recognizing that quantifying matches differs from quantifying appearance. Color matching tells us whether two lights match one another. If we place these lights in a different context, or change their shape, they continue to match but the appearance (of both) will change. Color matching was solved, even though the interesting problem of color appearance requires further investigation that accounts for context as well as stimulus pattern and dynamics.

### 4.4. Virtual color channels

The subspace-intersection idea (Section 2.2) may have value in modern image systems applications. One potential application is a new way to relate the responses of two cameras with different color channels, or one camera and the human visual encoding, or between two different life forms (person and pet; predator and prey).

The key idea is to create a virtual color channel by the intersection of the subspaces defined by the color channels in the two systems. We describe the virtual channel idea here because it differs from the conventional approach to matching data between two different systems.

#### 4.4.1. Low dimensional example

We visualize the virtual color channel concept in Figure 5, again showing the reduced case of three wavelengths to make the graphical representation straightforward. In this case, we illustrate two cameras, each with two color channels. The spectral quantum efficiency of each channel is specified by a 3-vector; the two vectors for cameras A and B, and the planes they span, are shown in Figure 5.

**Fig. 5.**
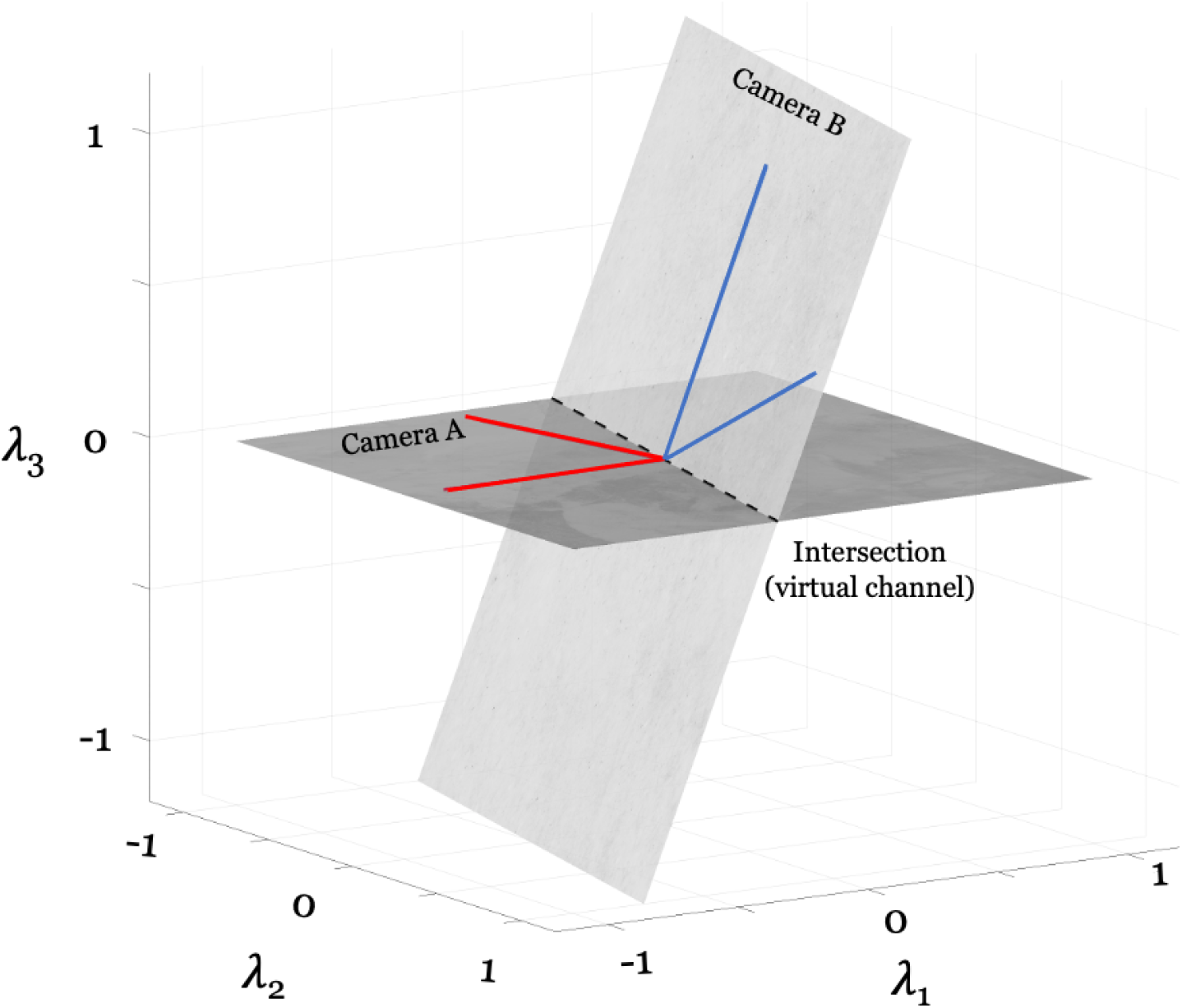
The virtual channel. In a world comprising three wavelengths, we can specify a color channel as a 3-vector. The red lines represent two color channels in camera A; the dark plane shows the range of channels that can be computed as a weighted sum of these two channels. The blue lines show two channels in camera B. No linear transformation of the camera A channels closely approximates the camera B channels. The light plane shows the range of channels that can be computed for camera B. The intersection of the two planes (dashed line) is a virtual channel that is in common between cameras A and B. The image contrast from the two cameras can be compared quantitatively on this virtual channel. See s_cfVirtualChannels.m.

Current practice relates data from two cameras by finding a linear transform that maps camera A channels into close approximations of camera B channels [42–44]. The range of possible channels that can be created from camera A falls within the dark-shaded plane in Figure 5. The range of linear combinations of the camera-B channels is shown by the light-shaded plane. The two camera-B channels are far from the plane of possible camera-A channels. Hence, the best approximation to the camera-B channels will be poor.

The intersection of the two planes is a virtual channel available to both cameras, but present in neither. The cameras have no channels in common, and no linear combination of channels in either camera matches a channel in the other. But the virtual channel is available to both and can be used to meaningfully compare image contrast for any light.

#### 4.4.2. Higher dimensional example

In the example above, we illustrated the case of two cameras with two channels for didactic purposes. As we consider the more realistic case of higher dimensional spectra, and different numbers of cameras and channels, there are other possibilities. Consider the case of two cameras, each with three spectral channels. We represent the channel spectral quantum efficiencies of the two cameras in the columns of two matrices **C** = [**r, g, b**] and 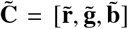. We use the intersection method to identify virtual color channels that are available to both cameras (Section 2.2).

To find the virtual channels we solve for two 3-dimensional column vectors, **v** and 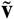, such that 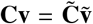. As before, we create a matrix that joins the six color channels in its columns, 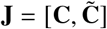and analyze the null space of this matrix. There are four possibilities corresponding to the plausible ranks of the matrix, **J**. The camera color matrices, **C** and 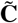

- rank 3: The camera channels occupy the same three-dimensional space
- rank 4: The virtual channels span a plane
- rank 5: A single virtual channel
- rank 6: No commonality

Nearly universally, modern image systems relies on the rank 3 approximation [42–44]: there is a 3 × 3 matrix transformation that maps a vector measured by **C** into an estimate of the measurement that would have been obtained from 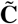. The virtual channel analysis provides a complementary approach, providing a method when **J** has rank 4 or 5. Figure 5 illustrates that a virtual channel may exist even though the 3 × 3 approximation is poor.

The brief development here is for a common case: two cameras each with three channels. There is a general formulation for *D* number of cameras, each with *N*_*D*_ channels, and there are multiple ways in which the virtual channel concept might be applied to complement the traditional analyses. We will treat this approach more fully in a future paper.

## 5. Conclusions

Color science has established a set of principles and standardized certain measurements that define how spectral electromagnetic radiation is encoded by the cone receptors in the human eye. This work was built on quantitative measurements using secure behavioral methods (matching). The principles established by Young in 1802 and the measurements reported by Maxwell in 1860 remain relevant today. Their work serves as the basis for all modern color technologies, including displays, cameras and printing.

Over the centuries, the mathematics underpinning these measurements has evolved. Young’s idea of a biological substrate was expressed and tested by Maxwell using a set of linear equations. His work took place at the time that Grassmann was developing a general mathematics of linear algebra and higher dimensional vector spaces [45], and these techniques have become a powerful tool in science and engineering. These tools now permeate the field of color science, particularly in the last few decades. This paper introduces new ideas based on this linear algebraic formulation.

First, we describe a novel method to solve the classic problem of identifying the cone fundamentals from the data of reduction dichromats. Our interest in this method arises because it uses only color matching functions from dichromats, requiring no auxiliary data, and can be applied simply without knowledge of or explicit correction for different sets of primaries used to collect data from different types of dichromats. The method calculates the intersection of the subspaces defined by the color matching functions of pairs of dichromats. We show that fundamentals derived using average color matching data from three types of dichromats are in good agreement with the most precise modern estimates of the cone fundamentals.

The estimates of cone fundamentals we provide here - based on these historical data - do not improve on the modern standard. Our interest is to describe the subspace intersection method; we analyze the historical data only to show the feasibility of the approach. By using the linear algebra framing, rather the specialized language of color science, we can generalize the method to a new image systems application. Specifically, the conventional approach for relating the images between two different color cameras, or between a camera and human visual system, is to find a linear transformation between the sensor encoding [42–44]. The subspace-intersection method generalizes this approach, explaining how to search for two linear transformations - one applied to each camera - that define a virtual channel where data from the two cameras can be compared quantitatively.

The subspace-intersection method simplifies the classic calculation of cone fundamentals in the sense that all of the data can be obtained using instrumentation with only two primary lights; there is no need to present three primaries and measure confusion lines. In addition, the method leads to the idea of a virtual channel, which may find applications in imaging systems, robotics and computer vision.

## Acknowledgments

To be added after review.

## Footnotes

1 The software repository that accompanies this paper includes data and code that we used to convert the tables in Maxwell’s 1860 paper into the format of modern colorimetry and that produces Figure 1. At a time when reliability and replication of results is of great concern to all scientists, we are pleased to confirm Judd and report that Maxwell’s data, measured 160 years ago on two typical trichromats, are consistent with the modern standards.

2 During the double-blind review process, please contact the authors for a link to the repository. Reviewers can make this request to the journal editor.

3 Often, the confusion lines are plotted in chromaticity, and the intersection of confusion lines originating at different chromaticities is referred to as the co-punctal point. Co-punctal points may also be used to define stimuli that isolate the class of cones that a dichromat is missing. We also note that confusion lines in the trichromatic space can be derived from the color matching functions of the dichromats.

4” … modern tendency in sensory physiology, which has found its most acute expression in the Physiological Optics of Helmholtz, is not leading us to the truth, and whoever wishes to open up new avenues of research in this area, must first free himself from the theories which now prevail (E. Hering, quoted in [41]).

## References

1. J. C. Maxwell, “IV. on the theory of compound colours, and the relations of the colours of the spectrum,” Philos. Trans. R. Soc. Lond. 150, 57–84 (1860).

2. J. D. Mollon, “The origins of modern color science,” in The Science of Color, S. K. Shevell, ed. (Elsevier, 2003), pp. 1–39.

3. T. Young, “On the theory of light and colours,” Philos. Trans. Royal Soc. Lond. 92, 20–71 (1802).

4. J. C. Maxwell, “XVIII.—Experiments on colour, as perceived by the eye, with remarks on Colour-Blindness,” Earth Environ. Sci. Trans. R. Soc. Edinb. 21, 275–298 (1857).

5. J. C. Maxwell, “The diagram of colors,” Trans. R. Soc. Edinb. 21, 275–298 (1857).

6. D. B. Judd, “Maxwell and modern colorimetry,” The J. Photogr. Sci. 9, 341–352 (1961).

7. D. B. Judd, “Fundamental studies of color vision from 1860 to 1960,” Proc. Natl. Acad. Sci. U. S. A. 55, 1313–1330 (1966).

8. J. C. Maxwell and Q. Zaidi, “On the theory of compound colours, and the relations of the colours of the spectrum,” Color. Res. Appl. 18, 270–287 (1993).

9. H. Grassmann et al., “On the theory of compound colors,” Phil. Mag 7, 254–64 (1854).

10. “James clerk maxwell produces the first color photograph,” https://www.historyofinformation.com/detail.php?id=3666. Accessed: 2023-8-15.

11. A. Stockman and L. Sharpe, “Spectral sensitivities of the middle-and long-wavelength sensitive cones derived from measurements in observers of known genotype,” Vis. Res. 40, 1711–1737 (2000).

12. A. Stockman, “Cone fundamentals and CIE standards,” Curr. Opin. Behav. Sci. 30, 87–93 (2019).

13. D. Brewster, “IV. on a new analysis of solar light, indicating three primary colours, forming coincident spectra of equal length,” Earth Environ. Sci. Trans. R. Soc. Edinb. 12, 123–136 (1834).

14. H. Helmholtz, “LXXXI. on the theory of compound colours,” The London, Edinburgh, Dublin Philos. Mag. J. Sci. 4, 519–534 (1852).

15. R. S. Turner, “The origins of colorimetry: What did helmholtz and maxwell learn from grassmann?” in Hermann Günther Graßmann (1809–1877): Visionary Mathematician, Scientist and Neohumanist Scholar, (Springer, 1996), pp. 71–86.

16. R. Heesen, “The Young-(Helmholtz)-Maxwell theory of color vision,” (2015).

17. B. A. Wandell, Foundations of Vision (Sinauer, Sunderland, MA, 1995).

18. A. Konig and C. Dieterici, “Die grundempfindungen und ihre Intensitäts-Vertheilung im spectrum (fundamental sensations and their intensity distribution in the spectrum),” Sitzungsberichte der Akademie der Wissenschaften pp. 805–829 (1886).

19. P. J. Bouma, “Mathematical relationship between the colour vision systems of trichromats and dichromats,” Physica 9, 773–784 (1942).

20. D. B. Judd, “Standard response functions for protanopic and deuteranopic vision,” J. Opt. Soc. Am. 35, 199 (1945).

21. N. D. Nuberg and E. N. Yustova, “Issledovanie cvetovogo zrenija dikhromatov,” Trudy Gosudarstvennogo Opt. Instituta (1955).

22. D. B. Judd, “Relation between normal trichromatic vision and dichromatic vision representing a reduced form of normal vision,” Acta Chromatica 1, 524–527 (1964). Available in NBS Special Publication Series 300, 1972, Precision Measurement and Calibration: Colorimetry, Volume 9, 147–150.

23. J. J. Vos and P. L. Walraven, “On the derivation of the foveal receptor primaries,” Vis. Res. 11, 799–818 (1971).

24. V. C. Smith and J. Pokorny, “Spectral sensitivity of color-blind observers and the cone photopigments,” Vis. Res. 12, 2059–2071 (1972).

25. CIE, “Fundamental chromaticity diagram with physiological axes – parts 1 and 2,” Tech. Rep. 170–1, Central Bureau of the Commission Internationale de l’Éclairage, Vienna (2007).

26. J. J. Vos, O. Estévez, and P. L. Walraven, “Improved color fundamentals offer a new view on photometric additivity,” Vis. Res. 30, 937–943 (1990).

27. A. Stockman, D. I. MacLeod, and N. E. Johnson, “Spectral sensitivities of the human cones,” J. Opt. Soc. Am. A Opt. Image Sci. Vis. 10, 2491–2521 (1993).

28. A. Stockman, “Colour and vision research laboratory,” http://www.cvrl.org/ (2001). Accessed: 2019-2-19.

29. M. A. Webster and D. I. A. MacLeod, “Factors underlying individual differences in the color matches of normal observers,” J. Opt. Soc. Am. A 5, 1722–1735 (1988).

30. Y. Asano, M. D. Fairchild, and L. Blonde, “Individual colorimetric observer model,” PLoS ONE 11, e0145671 (2016).

31. A. Stockman and A. T. Rider, “Formulae for generating standard and individual human cone spectral sensitivities,” Color. Res. & Appl. 48, 818–840 (2023).

32. B. A. Wandell, D. H. Brainard, and N. P. Cottaris, “Visual encoding: Principles and software,” Prog. Brain Res. 273, 199–229 (2022).

33. G. Strang, Introduction to linear algebra (SIAM, 2022).

34. G. Wyszecki and W. Stiles, Color Science: Concepts and Methods, Quantitative Data and Formulae (John Wiley & Sons, New York, 1982), 2nd ed.

35. H. Moreira, L. Álvaro, A. Melnikova, and J. Lillo, Colorimetry and Dichromatic Vision (InTech, 2018).

36. F. Pitt, “Characteristics of dichromatic vision,” in Reports of the Committee upon the Physiology of Vision, (Medical Research Council, 1935), 14, pp. 1–55.

37. W. D. Wright, Colour Vision. Research on Normal and Defective Colour Vision (C.V. Mosby, 1938).

38. W. D. Wright, “The characteristics of tritanopia,” J. Opt. Soc. Am. 42, 509–521 (1952).

39. D. A. Baylor, “Photoreceptor signals and vision. proctor lecture,” Invest. Ophthalmol. Vis. Sci. 28, 34–49 (1987).

40. E. Hering, Grundzüge der Lehre vom Lichtsinn (1874) (Outlines of a theory of the light sense, translated by Hurvich and Jameson) (Harvard University Press, Cambridge, MA, 1964).

41. I. P. Howard, “The Helmholtz–Hering debate in retrospect,” Perception 28, 543–549 (1999).

42. M. S. Brown, “Color processing for digital cameras,” in Fundamentals and Applications of Colour Engineering, P. Green, ed. (Wiley, 2023), pp. 81–98.

43. M. Delbracio, D. Kelly, M. S. Brown, and P. Milanfar, “Mobile computational photography: A tour,” Annu. Rev Vis Sci 7, 571–604 (2021).

44. M. Afifi and M. S. Brown, “Sensor-independent illumination estimation for dnn models,” arXiv preprint arXiv:1912.06888 (2019).

45. H. Grassmann, “Zur Theorie der Farbenmischung,” Ann. der Physik und Chemie 165, 69–84 (1853).

